# Highly specific chimeric DNA-RNA guided genome editing with enhanced CRISPR-Cas12a system

**DOI:** 10.1101/2021.09.04.458978

**Authors:** Hanseop Kim, Wi-jae Lee, Chan Hyoung Kim, Yeounsun Oh, Lee Wha Gwon, Hyomin Lee, WooJeung Song, Junho K. Hur, Kyung-Seob Lim, Kang Jin Jeong, Ki-Hoan Nam, Young-Suk Won, Youngjeon Lee, Young-Hyun Kim, Jae-Won Huh, Bong-Hyun Jun, Dong-Seok Lee, Seung Hwan Lee

## Abstract

The clustered regularly interspaced short palindromic repeats (CRISPR)-Cas12a system is composed of a Cas12a effector that acts as a deoxyribonucleic acid (DNA)-cleaving endonuclease and a crispr ribonucleic acid (crRNA) that guides the effector to the target DNA. It is considered a key molecule for inducing target-specific gene editing in various living systems. Here, we improved the efficiency and specificity of the CRISPR-Cas12a system through protein and crRNA engineering. In particular, to optimize the CRISPR-Cas12a system at the molecular level, we used a chimeric DNA-RNA guide chemically similar to crRNA to maximize target sequence specificity. Compared to the wild type (wt)-Cas12a system, when using enhanced Cas12a system (en-Cas12a), the efficiency and target specificity improved on average by 7.41 and 7.60 times respectively. In our study, when the chimeric DNA-RNA guided en-Cas12a effector was used, the gene editing efficiency and accuracy were simultaneously increased. These findings could contribute to highly accurate genome editing, such as human gene therapy, in the near future.

## Introduction

The clustered regularly interspaced short palindromic repeats (CRISPR)-Cas system, which is known to be a bacterial defense system, is composed of Cas endonuclease and guide ribonucleic acid (RNA); it is known to operate in various living organisms (1–3). Recently, it has been used as a key tool for *in vivo* therapeutics because it can be reprogrammed specifically for a target gene. Thus, it is easy to use the CRISPR-Cas system to access genetic diseases (4, 5). The field of gene therapy is growing into a large market in which these advanced genome editing tools are frequently employed; thus, it is important to determine whether the CRISPR-Cas system can accurately induce mutations into a target (6, 7). The target sequence specificity of CRISPR occurs due to molecular-level interactions resulting from its intrinsic properties (8–10). CRISPR-Cas endonuclease recognizes target deoxyribonucleic acid (DNA) based on the complementary nucleotide sequence contained in the guide RNA. CRISPR-Cas recognizes the protospacer adjacent motif (PAM) sequence in the target gene through the PAM interaction (PI) domain, melts the DNA double helix, and propagates the hybridization of guide RNA and target DNA to form a stable R-loop that induces target DNA cleavage (11–14). It has been reported that the hybridization between the guide RNA and the target DNA, which is formed to aid the CRISPR-Cas system in stably binding to the target DNA, is approximately 20-24 bp; it can have various mismatch tolerances depending on the target sequence (15–19). Accordingly, the possibility of inducing cleavage to off-targets similar to the target sequence has been reported, and efforts have been made to reduce such errors (9, 20–22).

Among CRISPR-Cas endonucleases, the CRISPR-Cas12a system, which belongs to Class II and type V, has excellent target specificity. Therefore, it has attracted much attention as an accurate genome editing tool for use as a therapeutic agent for human beings in the future (14, 23–26). Unfortunately, the CRISPR-Cas12a system has also been reported to have a tolerance for mismatches in the intermediate region (8-9 bp), or in the PAM distal region inside the protospacer, which is necessary for target recognition (16). This off-target cleavage effect appears to be more serious for engineered CRISPR-Cas12a, which has enhanced target recognition and improved gene editing efficiency (27, 28). When considering gene therapy for human systems in the future, efficiency and safety will likely be important issues; they must be addressed simultaneously to improve the CRISPR-Cas12a system.

In this study, we devised a technology that dramatically lowers the induction of off-target mutations, while efficiently inducing on-target mutations by effectively recognizing various target nucleotide sequences in human-derived cell lines. When using this enhanced Cas12a system (en-Cas12a) with strong target recognition, the average efficiency of inducing mutations in the target sequence increased (1.7-17.16 fold) when guided by a chimeric DNA-RNA guide, compared to the wild-type Cas12a (wt-Cas12a) system. In addition, the average (0.5-10.6%) of the mutation induction efficiency of off-target nucleotide sequences was reduced (0.1-3.6%) by using a chimeric DNA-RNA guide, which increased the target specificity 7.6-fold on average. Using the chimeric DNA-RNA guide-based en-Cas12a system developed in this study, it is possible to induce target-specific, high-efficiency gene editing. Therefore, our proof of concept study could contribute to the fundamental treatment of various incurable human diseases resulting from genetic mutations in the near future.

## Results

### Comparison of target DNA cleavage activity of chimeric DNA-RNA-guided engineered en-AsCas12a and wt-AsCas12a

The CRISPR-Cas12a system uses single-stranded crispr RNA (crRNA) to hybridize target DNA with 20 bases, form a stable R-loop, and induce target DNA cleavage. When the amino acid residues (Lys548, Ser542, and Glu174) interacting near the PAM (TTTN) sequence were changed to positive residues for the interaction between Cas12a and the target DNA **(Fig. 1a, left inset)**, the target-induced indel ratio (%) was improved for various genes (28). From these results, we speculate that PAM recognition contributes to the kinetics of the entire Cas12a target recognition and DNA cleavage process, and that it can eventually affect stable R-loop formation through the hybridization of DNA and crRNA. The target specificity (on-target editing/off-target editing) of the Cas12a system has previously been optimized by substituting DNA for the 3’-end of the crRNA to change the hybridization energy between crRNA and target DNA **(Fig. 1a, right inset)** (29). Based on this system, here we attempted to maximize the target specificity and genome editing efficiency using en-AsCas12a(*Acidaminococcus* sp. Cas12a), which has enhanced target recognition. First, to improve target specificity by changing the binding energy of the target DNA-crRNA hybridization region, we gradually substituted the crRNA with DNA; we then confirmed the influence of this substitution on the target DNA cleavage for wt-AsCas12a and en-AsCas12a effectors **(Fig. 1b)**. Amplicon cleavage assays were performed on the target nucleotide sequences of both genes (*DNMT1* and *CCR5* site2) **(Supplementary Fig. 1)**. When DNA was gradually substituted from the 3’-end of the crRNA (recognized by AsCas12a), en-AsCas12a showed improved target recognition compared with wt-AsCas12a; it demonstrated more tolerance to the DNA substitution of crRNA **(Fig. 1b, Supplementary Fig. 2)**. As previously reported (29), when 12 or more DNAs are substituted from the 3’-end of Cas12a, the cleavage activity of the DNA amplicon is reduced, and almost no activity is shown in substitutions over 16 nt DNAs. However, en-AsCas12a showed robust cleavage activity after 12 nt DNA substitutions at the 3’-end of crRNA, but showed a decrease in activity by more than half following 16 nt or more DNA substitutions. This indicates that en-AsCas12a is more tolerant to DNA substitution at the 3’-end of crRNA than wt-AsCas12a, which is advantageous for target DNA cleavage.

**Figure 1.**
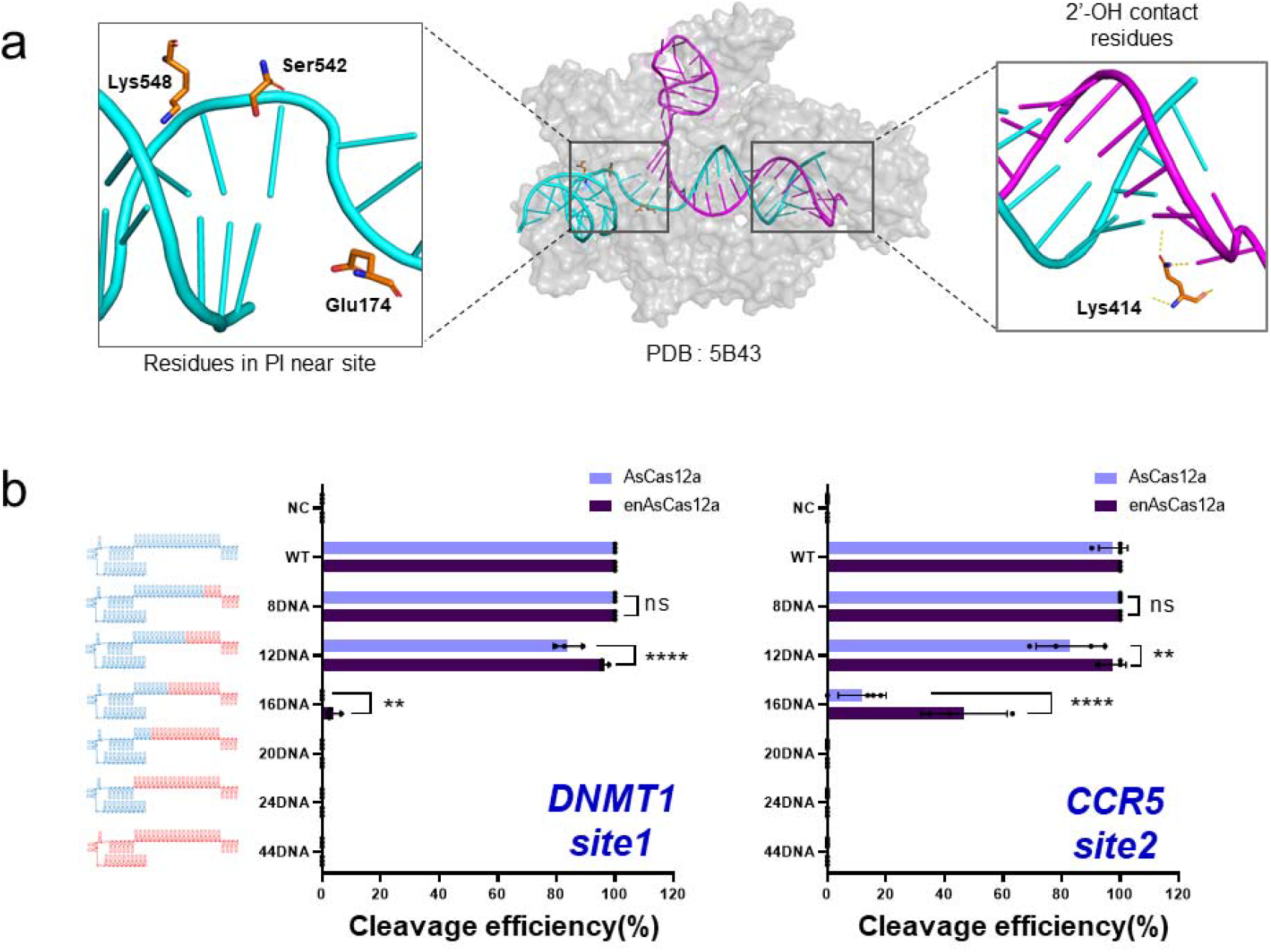
Comparison of target DNA cleavage activity of chimeric DNA-RNA guided en-Cas12a and wt-Cas12a. a, Structure of target-strand DNA-crRNA-AsCas12a complex(PDB: 5B43). left inset: amino acids in AsCas12a interacting with around the PAM sequence in the target DNA, right inset: Amino acid (Lys414) interacting with the target-strand DNA-crRNA duplex in AsCas12a (hydrogen bonding with the 2’-OH group on the crRNA 3’-end side). b, Comparison of cleavage efficiency of wt-AsCas12a and en-AsCas12a using a chimeric DNA-RNA guide. Comparison of cleavage efficiency of DNA amplicons including target nucleotide sequences (*DNMT1, CCR5-site2*) with gradual DNA substitution from the 3’-end of crRNA. NC: Negative control, WT: Wild-type crRNA was treated with wt- or en-AsCas12a. The RNA region of the (cr)RNA is shown in blue, and the substituted DNA region is shown in red (‘8-44DNA’ indicates a number of substituted DNA nucleotides in (cr)RNA). The X-axis indicates the efficiency of the target gene (*DNMT1, CCR5*) cleavage by wt- and en-AsCas12a using various chimeric DNA-RNA guides (DNA substitution from the 3′-end of the (cr)RNA). All cleavage efficiency were calculated from agarose gel separated band intensity (cleaved fragment intensity (%) / total fragment intensity (%)) and normalized to wild-type (cr)RNA **(Supplementary Fig. 2)**. Data are shown as means ± s.e.m. from three independent experiments. *P*-values are calculated using a two-way ANOVA with sidak’s multiple comparisons test (ns: not significant, P*:<0.0332, P**:<0.0021, P***:<0.0002, P****:<0.0001).

### Optimization of the genome editing activity of engineered en-AsCas12a based on a chimeric DNA-RNA guide to a target nucleotide sequence on the intracellular genome

To check whether the engineered en-AsCas12a effector, based on this chimeric DNA-RNA guide, could effectively induce target-specific gene mutations in human cells, various chimeric DNA-RNA guides were used to induce mutations and the efficiency was analyzed in comparison with wt-AsCas12a. Comparative analysis of mutation induction efficiencies for the target nucleotide sequences of three genes (*DNMT1, IL2A-AS1*, and *CCR5 site1*), revealed that engineered en-AsCas12a outperformed wt-AsCas12a in terms of editing efficiency **(Fig. 2, Supplementary Fig. 1)**. In particular, the induction of gene mutations targeting intracellular loci showed a different trend from that of amplicon cleavage **(Fig. 1b)**. Unlike wt-AsCas12a, which exhibited a significantly lower operating efficiency based on chimeric DNA-RNA guides for the target sequence in the genome, engineered en-AsCas12a allowed up to 8 nt DNA substitutions (8DNA) from the 3’-end of the crRNA while maintaining the editing activity **(Fig. 2, Supplementary Fig. 3)**. Surprisingly, engineered en-AsCas12a showed 1.5-to 13.6-fold improvements in mutation induction efficiency compared to wt-AsCas12a when the 3’-end of crRNA was substituted with 8 nt DNA to increase target specificity. This effect was universally confirmed in various genes (*CCR5 site2, FANCF*); on average, a 7.3-fold higher recovery was achieved **(Supplementary Figs. 3, 4)**. Previous studies have reported that when AsCas12a targets the intracellular genome and operates based on a chimeric DNA-RNA guide, it is difficult to induce mutations in the target nucleotide sequence due to many restrictions on the topology of the intracellular genome (29). Accordingly, in this study, we attempted to change the genome topology near the target sequence by using nickase. We compared the resulting changes in the genome editing efficiency of the target sequence for wt-AsCas12a and en-AsCas12a **(Supplementary Figs. 1, 3, 4)**. In the case of wt-AsCas12a, the mutation induction efficiency, which was reduced by the use of a 8 nt DNA substituted chimeric DNA-RNA guide (8DNA), was completely recovered by co-treatment with nickase. In the case of en-AsCas12a, the mutation induction efficiency was maintained at a level similar to that of wt-crRNA using chimeric DNA-RNA (8DNA); it was not significantly affected by nickase **(Fig. 2, Supplementary Figs. 3, 4)**. Therefore, these data indicate that using the en-AsCas12a effector, based on the chimeric DNA-RNA guide (8DNA), enables more effective genome editing than wt-AsCas12a when inducing mutations on the target DNA, without the help of nickase.

**Figure 2.**
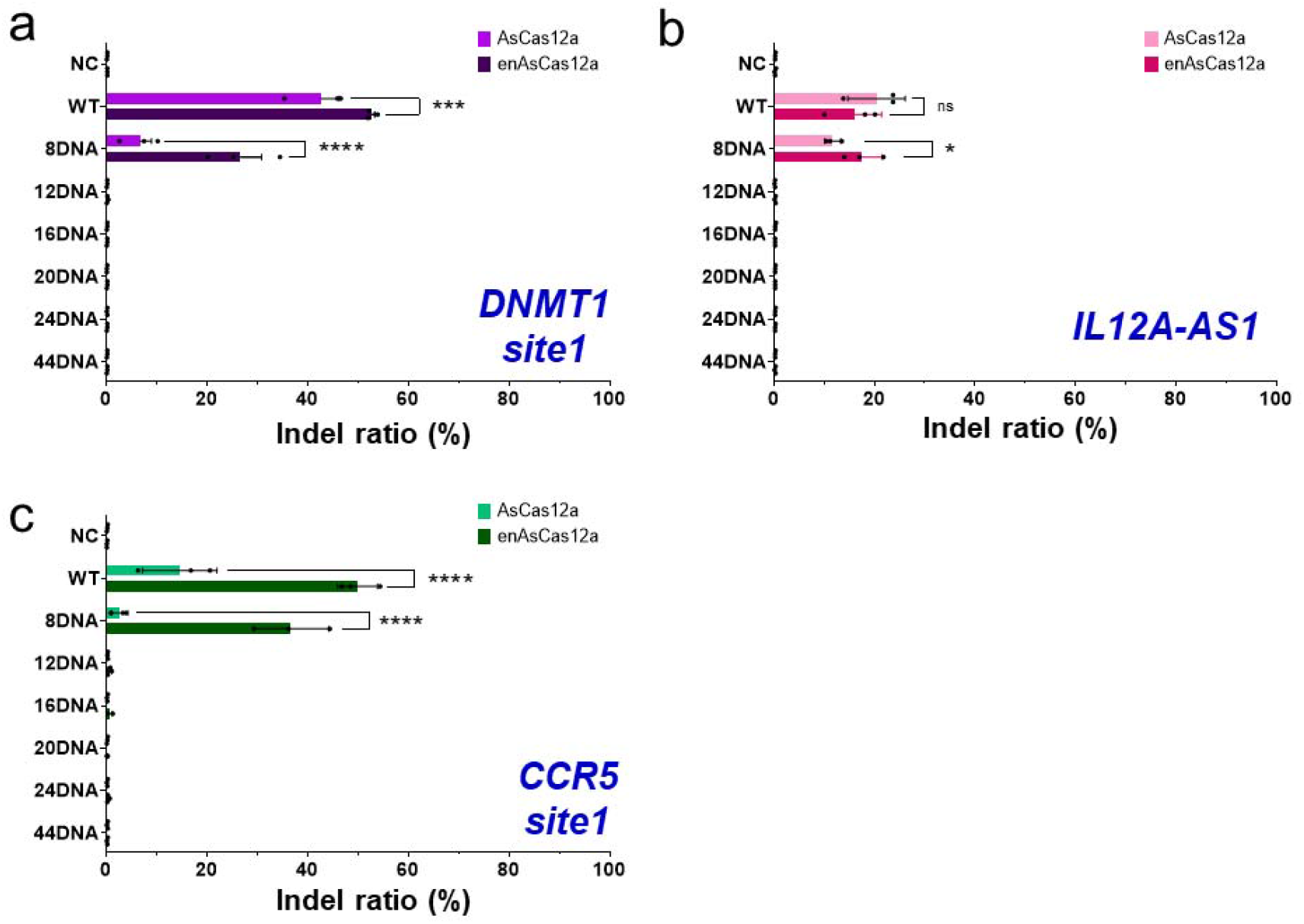
Comparison and optimization of genome editing efficiency of en-Cas12a and wt-Cas12a based on chimeric DNA-RNA guide targeting intracellular genome. a-c. Comparison of genome editing efficiency (%) of wt-AsCas12a and en-AsCas12a using a chimeric DNA-RNA guide for human-derived cell line (HEK293FT). Comparison of indel induction efficiency (%) in intracellular genome target sequences (*DNMT1, IL2A-AS1, CCR5-site1*) by gradual DNA substitution of the 3’-end of crRNA. All the indel ratio was calculated from targeted amplicon sequencing (Indel ratio(%) = mutant DNA read number / total DNA read number). Data are shown as means ± s.e.m. from three independent experiments. *P*-values are calculated using a two-way ANOVA with sidak’s multiple comparisons test (ns: not significant, P*:<0.0332, P**:<0.0021, P***:<0.0002, P****:<0.0001). NC: negative control, WT: Wild-type crRNA was treated with wt- or en-AsCas12a, 8-44DNA: Chimeric crRNA (sequential 8-44DNA substitution at 3’-end of crRNA) was treated with wt- or en-AsCas12a.

### Improving target specificity for inducing genetic mutations in the intracellular genome using chimeric DNA-RNA guide-based engineered en-AsCas12a

Next, we compared the chimeric DNA-RNA guide-based engineered en-AsCas12a and wt-AsCas12a effectors regarding their target specificities under optimized conditions (3’-end 8DNA substitution of crRNA) that effectively induced mutations in the target sequence **(Fig. 3, Supplementary Fig. 1)**. In the case of wt-Cas12a-based targeting of the *CCR5-site1*, the target mutation induction efficiency was greatly reduced by the use of chimeric DNA-RNA **(Fig. 3a)**. Unlike wt-AsCas12a, in which mutation induction efficiency was recovered in a nickase-dependent manner, for en-AsCas12a the target mutation efficiency was maintained in a nickase-independent manner by using chimeric DNA-RNA, in which the 3’-end was substituted with 8 nt DNA **(Fig. 3b)**. The nickase dependency was lowered by 11.25 times **(Fig. 3c)**, and target specificity was increased by 2.79 times **(Fig. 3d)** using en-AsCas12a. In addition, when the chromosome topology near the target sequence was changed using nickase, the target specificity using chimeric DNA-RNA was further increased by 3.45-fold, compared to that of wt-AsCas12a **(Fig. 3d)**. We further compared the target specificities of the engineered en-AsCas12a and wt-AsCas12a effectors in the target sequences of two other genes (*AAVS1-site1* and *DNMT1-site2*) **(Fig. 3e-l)**. When a chimeric DNA-RNA guide was used for *AAVS1-site1*, neither en-AsCas12a nor wt-AsCas12a effectors showed nickase dependence, and gene mutations were induced with similar efficiencies to conditions using wt-crRNA **(Fig. 3e-h)**. However, when using the en-AsCas12a effector to target the *AAVS1-site1* sequence, the indel ratio (%) was significantly higher (3.9-fold) than that of wt-Cas12a. However, there were also more unintentional mutations in the off-target sequence (off-target1) **(Fig. 3e, f)**. We confirmed that the number of mutations induced in the off-target1 sequence was dramatically reduced by the use of chimeric DNA-RNA, in which the 3’end was substituted with 8 nt DNA. Therefore, the overall target specificity was increased 3.5-fold compared to that of wt-AsCas12a when chimeric DNA-RNA (8DNA) guided en-AsCas12a was used **(Fig. 3h)**. The *DNMT1-site2* showed the same trend as *the AAVS1-site1* locus targeted by the Cas12a system **(Fig. 3i-l)**. In the case of the en-AsCas12a effector based on the chimeric DNA-RNA guide with 8 nt DNA substitution at the 3’-end, the indel ratio (%) was significantly increased (1.9-fold) compared to that of wt-AsCas12a. Furthermore, the off-target nucleotide sequence (off-target1, 2)-induced mutations were dramatically reduced **(Fig. 3i, j)**. As a result, the target specificity was increased twofold, regardless of the use of nickase **(Fig. 3k, l)**. In conclusion, by inducing mutations in three genes using chimeric DNA-RNA guided engineered en-AsCas12a, the targeted indel ratio (%) was improved 7.3-fold on average, without the help of nickase. The off-target mutation induction efficiency was also reduced. Eventually, the target specificity was improved 3.1-fold compared to that of wt-AsCas12a.

**Figure 3.**
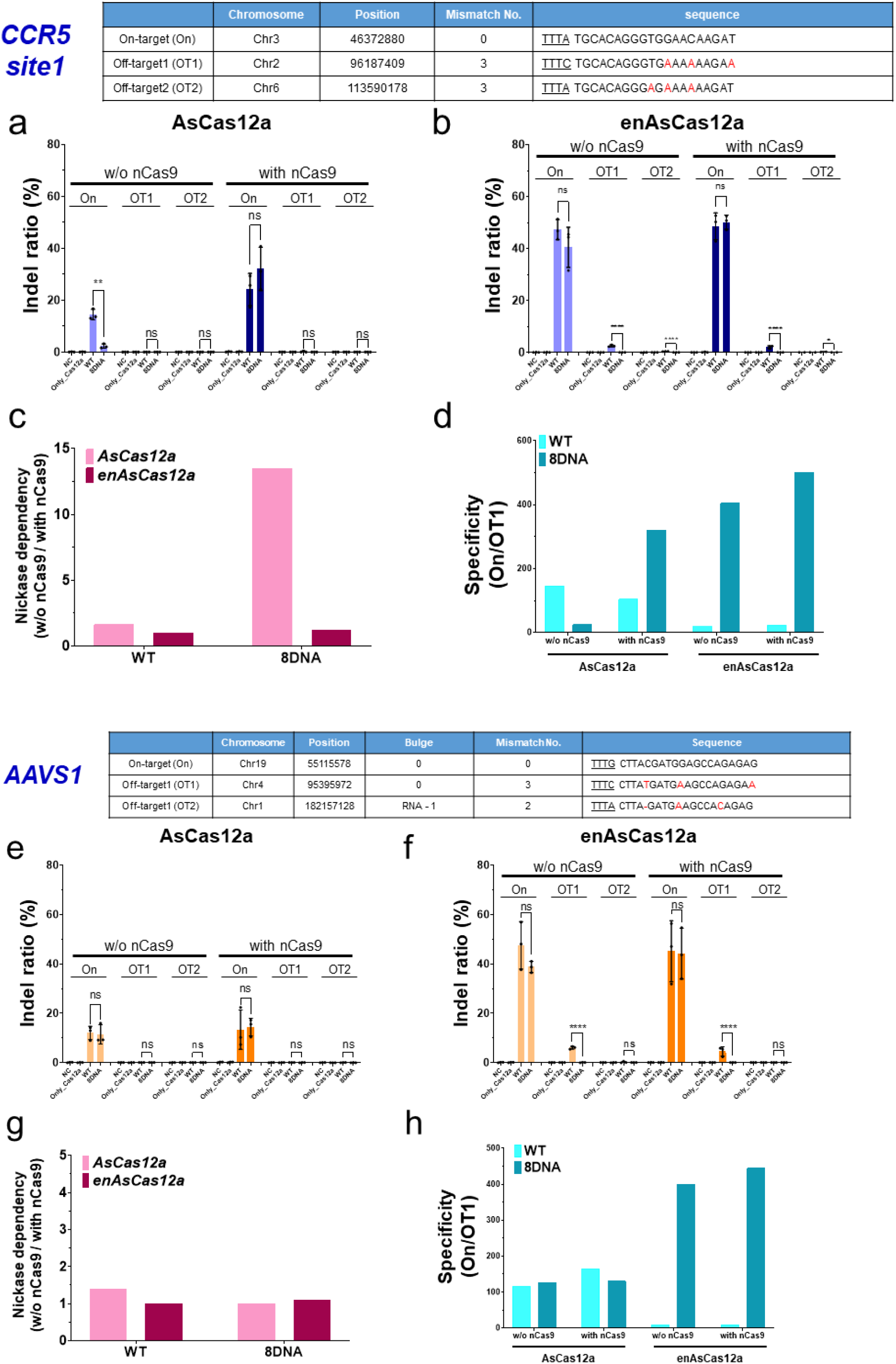
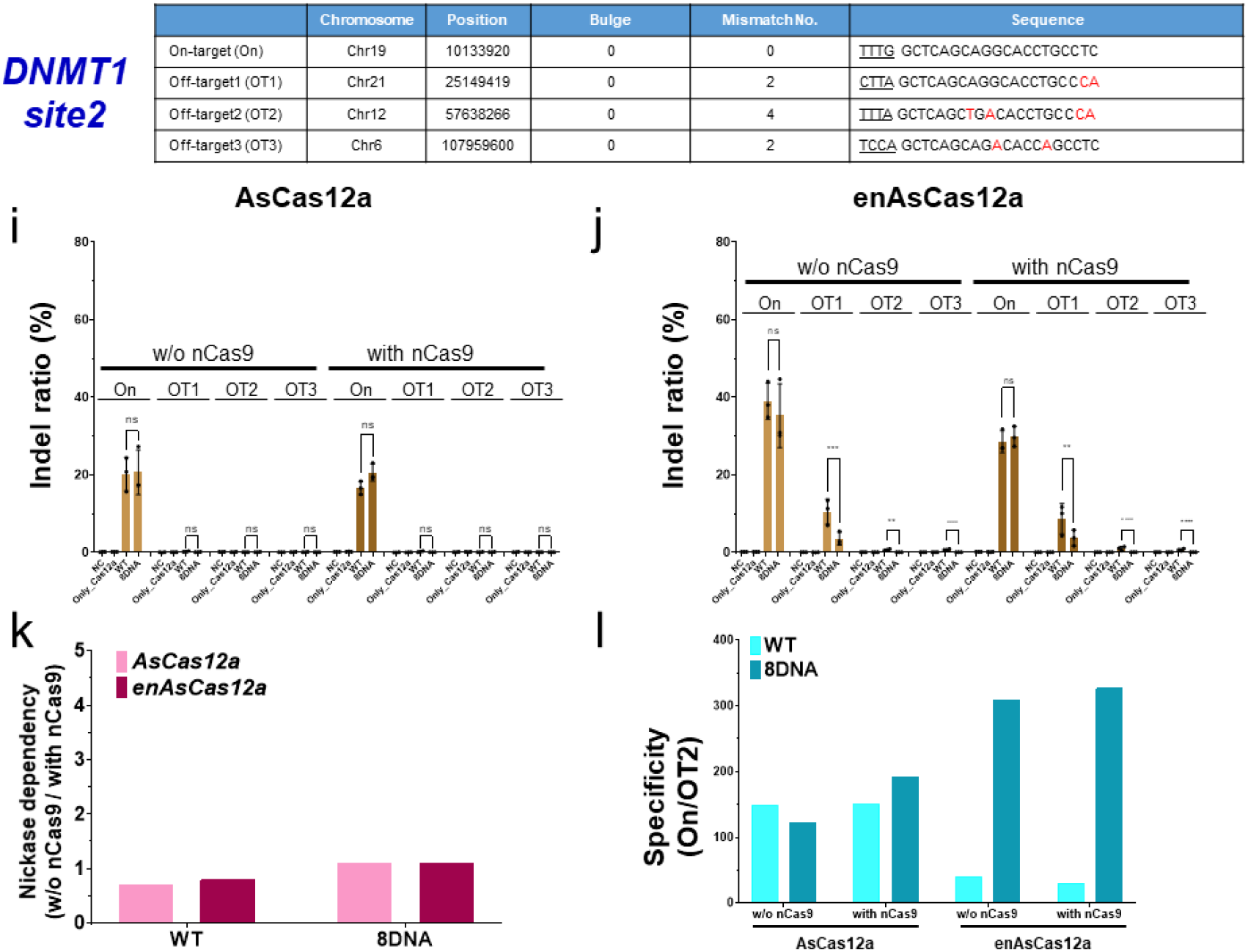
Comparison of genome editing specificity of en-AsCas12a and wt-AsCas12a based on chimeric DNA-RNA guide targeting intracellular genome. a-l, Comparison of genome editing target specificity (on-target editing(%) / off-target editing(%)) of wt-AsCas12a and en-AsCas12a on genome in human cell line(HEK293FT) using an optimized chimeric DNA-RNA guide(8DNA). a-d, Comparison of target specificity between wt- and en-AsCas12a on the intracellular genome target sequence (*CCR5-site1*) using wt-crRNA(WT) and 3’-end 8nt DNA substituted crRNA(8DNA). e-h, Comparison of target specificity on the target sequence (*AAVS1-site1*). i-l, Comparison of target specificity on the target sequence (*DNMT1-site2*). Data are shown as means ± s.e.m. from three independent experiments. *P*-values are calculated using a two-way ANOVA with sidak’s multiple comparisons test (ns: not significant, P*:<0.0332, P**:<0.0021, P***:<0.0002, P****:<0.0001). NC: negative control, only Cas12a: only protein treated, WT: Wild-type crRNA was treated with wt- or en-AsCas12a, 8DNA: Chimeric crRNA (sequential 8DNA substitution at 3’-end of crRNA) was treated with wt- or en-AsCas12a. nCas9: nickase Cas9(D10A), Nickase dependency = (w/o nCas9 editing(%) / w/ nCas9 editing(%)), Specificity = (on-target editing(%) / off-target editing(%)).

### A model for improving target specificity and mutation induction efficiency of en-Cas12a, based on chimeric DNA-RNA guides

Combining all of the above findings, the results of the working mechanism of en-Cas12a, compared to the existing wt-Cas12a, are presented **in Fig. 4**, based on the chimeric DNA-RNA guide. When wt-Cas12a was used to cleave the target sequence in the genome of the cell, there was tolerance for mismatches between the protospacer (20bp) middle part and the PAM (TTTN) distal region, so there was a possibility that Cas12a could recognize and cleave off-target sequences. When wt-Cas12a, based on the chimeric DNA-RNA guide, was used (in which the 3’-end was substituted with 8 DNAs), the off-target effect could be reduced by changing the binding energies of the target DNA and crRNA. However, when chimeric DNA-RNA based cleavage was performed on the genome in the cell, the mutation-inducing effect on the target nucleotide sequence was largely decreased. So, the target specificity was increased only by changing the topology of the genome with the simultaneous use of nickase.

**Figure 4.**
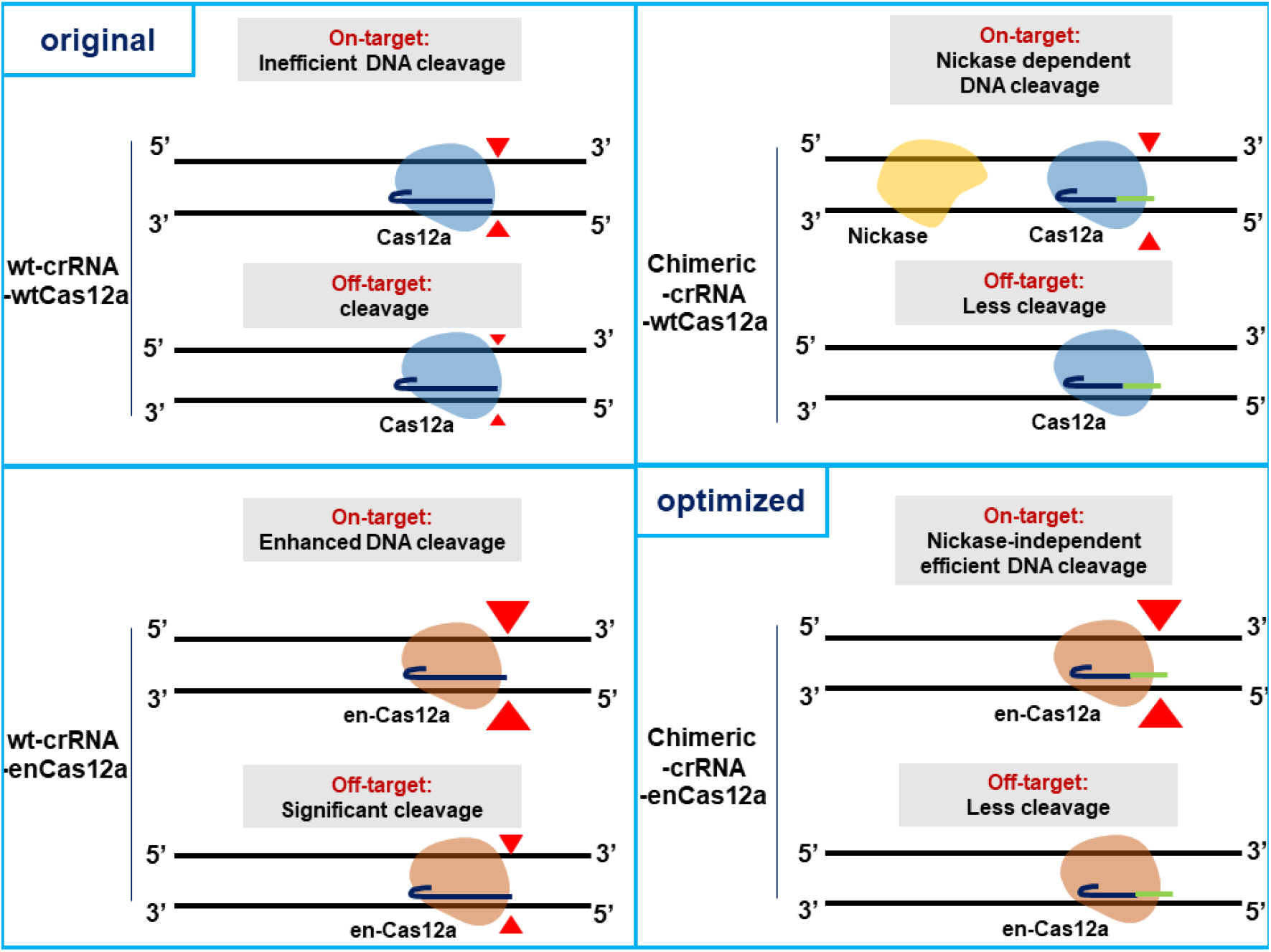
A model for enhancing target specificity and editing efficiency of en-Cas12a based on chimeric DNA-RNA guided engineering. wt-crRNA guided wt-Cas12a system: In general, the wt-Cas12a effector can induce genetic mutations in target sequences, but can also induce mutations in similar off-target sequences., wt-crRNA guided en-Cas12a system: Recognition of a target sequence is enhanced by a effector engineered by amino acid substitution, and thus the efficiency of inducing gene mutations is increased compared to that of the wt-Cas12a effector. However, due to the same principle as target sequence recognition, there is a problem in that the mutation induction efficiency of off-target sequences is also increased., chimeric-crRNA guided wt-Cas12a system: Effectively reduced off-target mutation induction efficiency when using a chimeric DNA-RNA guide with 8nt DNA substituted at the 3’-end. However, the efficiency of mutation induction for the target nucleotide sequence in the genome is also reduced, so the efficiency is restored only when there is the action of nickase on the nucleotide sequence near the target., chimeric-crRNA guided en-Cas12a system: Maximizes the target sequence indel ratio(%) and minimizes the off-target indel ratio(%) when using a chimeric DNA-RNA(8DNA) guided en-Cas12a effector which is engineered by amino acid substitution. It can induce more accurate and high-efficiency gene editing than wt-Cas12a on genomic DNA. DNA cleavage points are indicated by red arrows, and the degree of cleavage is indicated by arrow size according to the Cas12a activity. The wt-Cas12a and en-Cas12a effectors are shown in blue and brown, respectively. In the wt-crRNA and chimeric DNA-RNA guides, RNA is indicated in dark blue, and nucleotides replaced with DNA are indicated in green.

In the case of en-Cas12a, in which target sequence recognition is reinforced by engineering the interacting amino acids of the PAM sequence recognition part, its efficiency in inducing target sequence mutations for various genes was increased compared to that of Cas12a, but its unintended off-target effects also increased greatly. Using the chimeric DNA-RNA-based en-Cas12a effector to target the intracellular genome can dramatically reduce the effects of off-target mutations. Without the help of nickase, it is thus possible to increase the target sequence editing efficiency and dramatically lower the off-target sequence editing efficiency. Regarding the efficient induction of gene mutations with respect to the use of chimeric guides, accurate and high-efficiency gene targeting is possible when using chimeric DNA-RNA-based en-Cas12a.

## Discussion

The CRISPR-Cas12a effector is attracting attention as a potential future target-specific genome editing tool as it is known to be capable of inducing mutations in a target sequence on a desired gene; it also has the highest target specificity among previously known CRISPR systems. However, the Cas12a system has been reported to have lower activity than Cas9 in general, and there remains room to improve the properties of the endonuclease itself for applications in various *in vivo* conditions. Efforts have been made to more effectively recognize the target DNA sequence and induce cleavage by engineering the Cas12a system (27, 28). These studies have shown overall improved activity compared to wt-Cas12a in various genes, through enhanced binding, by changing amino acids around the domain, within Cas12a, and by recognizing the PAM sequence in the target DNA. However, most CRISPR endonucleases induce double-stranded cleavage by forming an R-loop, through the complementary binding of target-strand DNA and crRNA. In general, as the tolerance increases due to enhanced binding affinity, off-target binding also increases. Therefore, we believe that the effects of off-target binding would be maximized when using engineered Cas12a systems, so a method for increasing target specificity is also required, in parallel with methods to enhance activity.

Previous studies have improved target specificity by applying crRNA engineering to the Cas12a system; target-specific gene mutations have been effectively induced by optimizing the length of DNA substitution. Mismatch tolerance has been reduced by changing the complementary binding energy of target-strand DNA and crRNA through sequential 8nt DNA substitutions at the 3’-end of crRNA. This principle confirms that the induction of off-target mutations can be reduced while maintaining the efficiency of inducing target mutations. Based on this, here we used chimeric DNA-RNA crRNA to engineer en-AsCas12a, which displayed maximized target sequence recognition and improved mutation induction efficiency in various target nucleotide sequences, compared to wt-AsCas12a. Surprisingly, the induction of off-target mutations was dramatically decreased, and eventually the target specificity was largely improved. Interestingly, the observed discrepancy between amplicon cleavage **(Fig. 1b)** and genome editing inside the cell **(Fig. 2)** showed that the cleavage activity of Cas12a endonuclease is greatly influenced by DNA topology. Chimeric DNA-RNA crRNA-based en-AsCas12a and wt-AsCas12a displayed differing sensitivities to the intracellular genome, so en-AsCas12a showed a higher gene mutation induction efficiency than wt-AsCas12a. These results suggest that en-AsCas12a can, to some extent, overcome structures that are unfavorable to target DNA cleavage due to unstable R-loop formation (due to the DNA substitution of crRNA in the genomic sequence). This is achievable by enhancing PAM recognition through protein engineering. In general, the low operating efficiency induced in the intracellular genome by wt-Cas12a (based on the DNA substitution of crRNA to improve target specificity) could be recovered by changing the genome topology by using nickase around the target sequence. However, in the case of en-Cas12a, it was possible to induce original levels of mutation following up to eight DNA substitutions at the 3’ end of the crRNA, without the help of nickase. Therefore, when using chimeric DNA-RNA guide-based en-AsCas12a, it was possible to simultaneously improve the genome editing efficiency (%) and the target specificity (on-target editing [%]/off-target editing [%]) compared to those of wt-AsCas12a by changing the hybridization energy of the target DNA strand and crRNA.

In this study, we developed a technology that maximizes the safety and efficiency of genome editing using chimeric DNA-RNA crRNA-based en-Cas12a. In the future, many improvements are needed in terms of efficiency and safety regarding the application of gene therapy to humans using the CRISPR system, or for sophisticated gene editing in *in vivo* systems. Through this technology, it is expected that the safety and efficacy of various CRISPR endonucleases can be optimized in a similar way when applied *in vivo*.

## Methods

### Preparation of the CRISPR-Cas12a recombinant protein and chimeric guides

wt- and en-AsCas12a recombinant proteins were prepared for the *in vitro* DNA cleavage assay. Codon-optimized AsCas12a (*Acidaminococcus* sp. Cas12a) coding sequence was cloned into a pET28a bacterial expression vector and then transformed into BL21 (DE3) *Escherichia coli* cells. Transformed bacterial colonies were cultured at 37 °C until the optical density reached 0.6, after which isopropylthio-β-galactoside (IPTG) inoculation was performed. After 48 h, *E. coli* cells were precipitated at 4 °C and 5,000 rpm, following which the culture medium of the upper layer was removed. The precipitated *E. coli* cell pellet was resuspended in lysis buffer [10 mM β-mercaptoethanol, 300 mM NaCl, 20 mM Tris-HCl (pH8.0), 1 mM PMSF, and 1% TritonX-100]. In order to disturb the bacterial cell membrane, sonication was performed on ice water for 3 min, following which the cell lysate was precipitated for 10 min at 5,000 rpm at 4 °C. Next, the nitrilotriacetic acid (Ni-NTA) resin was pre-washed with wash buffer [20mM Tris-HCl (pH 8.0), 300 nM NaCl] and the precipitated cell lysate was stirred at 4 °C for 90 min. Washing was performed with ten times the volume of wash buffer to remove non-specific binding components in the mixed cell lysate solution. For the elution of AsCas12a protein, an elution buffer [20 mM Tris-HCl (pH 8.0), 300 nM NaCl, 200 mM imidazole] was used and finally exchanged against the storage buffer [200 mM NaCl, 50 mM 4-(2-hydroxyethyl)-1-piperazineethanesulfonic acid (HEPES; pH 7.5), 1 mM dithiothreitol (DTT), 40% glycerol] using a Centricon (Millipore, Amicon® Ultra-15), and stored at -80 °C. Chimeric DNA-RNA oligonucleotides (Bioneer) were synthesized for each target gene sequence and dissolved in diethyl pyrocarbonate (DEPC) water and then stored at -80 °C **(Supplementary Table S1)**.

### Preparation of the guide RNA for Cas12a and nCas9(D10A)

Guide RNAs for Cas12a and nCas9(D10A) were generated by *in vitro* transcription. A DNA template for *in vitro* transcription was constructed using annealing or extension polymerase chain reaction (PCR) with sense and antisense DNA oligonucleotides (Macrogen) containing the target DNA sequence **(Supplementary Table S2)**. DNA templates were mixed with T7 RNA polymerase (NEB, M0251L) and reaction buffer mixture (50 mM MgCl2, 100 mM ribonucleoside tri-phosphate (rNTP; Jena Bio, NU-1014), 10X RNA polymerase reaction buffer, 100 mM DTT, RNase inhibitor Murine, DEPC), and incubated at 37 °C. After 16 h, to remove the original DNA template, DNase I was added and the mixture was incubated for another 1 h at 37°C. The transcribed RNA was purified using a column (MP Biomedicals, GENECLEAN® Turbo Kit). The purified RNA was concentrated through lyophilization (2,000 rpm) at -55 °C for 1 h.

### *In vitro* DNA cleavage assay

On-/off-target site PCR amplicons of each gene (DNMT1, CCR5, IL2A-AS1, and AAVS1) were obtained from purified human genomic DNA using DNA primers **(Supplementary Table S2)**. The purified target PCR amplicon was incubated with purified recombinant wt- or en-Cas12a protein and crRNA (RNA-guides or chimeric DNA-RNA guides) in 10X buffer (NEBuffer3.1, NEB) for 1 h. After adding a stop buffer [100 mM ethylenediaminetetraacetic acid (EDTA), 1.2% sodium dodecyl sulfate (SDS)] to stop the reaction, the cleaved fragment was separated using 2% agarose gel electrophoresis. DNA cleavage efficiency (cleaved fragment intensity [%]/total fragment intensity [%]) was measured using ImageJ software (NIH).

### Cell culture and transfection

The HEK293FT (ATCC) cell line was cultured in Dulbecco’s modified Eagle medium (DMEM, Gibco) with 10% fetal bovine serum (FBS, Gibco) at 37 °C and in 5% CO_2_. To ensure efficient chimeric DNA-RNA guide delivery, we performed sequential transfection with wt- or en-Cas12a expression vectors and crRNAs. For the primary transfection, 10^5^ cells were mixed with a plasmid vector (AsCas12a, n-SpCas9 (D10A, H840A)) and 20 μl of electroporation buffer (Lonza, V4XC-2032) and were nucleofected according to the manufacturer’s instructions (program: CM-137). The transfected cells were transferred to a 24-well plate with 500 μl of media and incubated at 37 °C in 5% CO_2_. Twenty four hours after primary transfection, for secondary transfection, crRNA (200 pmol), single guide RNA (sgRNA; 30 pmol), 1 μl P3000, and 1.5 μl Lipofectamine 3000 reagent (Thermo) were mixed in 50 μl Opti-MEM (Gibco), incubated for 10 min, and added to DMEM media. Forty-eight hours after the second transfection, cells were harvested and genomic DNA was extracted using a genomic DNA purification kit (Qiagen, DNeasy Blood & Tissue Kit).

### Deep sequencing and data analysis

To analyze the indel frequency of the on-/off-target locus of each gene, targeted deep sequencing was performed using PCR amplicons. The Cas-OFFinder (http://www.rgenome.net/cas-offinder/) web tool was used to select potential off-target sites corresponding to each on-target site. For the preparation of PCR amplicons, PCR amplification was performed using DNA primers corresponding to each endogenous locus (**Supplementary Table S2**). To add adapter and index sequences to each 5’ and 3’ ends, nested PCR was performed using Phusion™ High-Fidelity DNA Polymerase (Thermo). After index tagging, the PCR amplicon mixture was analyzed using a Mini-Seq (Illumina, SY-420-1001) according to the manufacturer’s guidelines. Sequencing read fatstq files were analyzed using Cas-Analyzer (http://www.rgenome.net/cas-analyzer/), and the indel ratio (mutant DNA read number/total DNA read number) was calculated.

## AVAILABILITY

CRISPR RGEN Tools is an open-source collaborative initiative available in the repository (http://www.rgenome.net/).

## ACCESSION NUMBERS

Targeted deep sequencing data are available at the NCBI Sequence Read Archive (SRA) under the accession number SRP334002.

## SUPPLEMENTARY DATA

Supplementary Data are available at online.

## ACKNOWLEDGEMENT

The authors thank the members of the National Primate Research Center (NPRC) for their helpful discussions.

